# Involvement of the cerebellum in structural connectivity enhancement in episodic migraine

**DOI:** 10.1101/2024.04.26.591265

**Authors:** Ana Matoso, Ana R Fouto, Inês Esteves, Amparo Ruiz-Tagle, Gina Caetano, Nuno A da Silva, Pedro Vilela, Raquel Gil-Gouveia, Rita G Nunes, Patrícia Figueiredo

## Abstract

**Background:** The pathophysiology of migraine remains poorly understood, yet a growing number of studies have shown structural connectivity disruptions across large-scale brain networks. Although both structural and functional changes have been found in the cerebellum of migraine patients, the cerebellum has barely been assessed in previous structural connectivity studies of migraine. Our objective is to investigate the structural connectivity of the entire brain, including the cerebellum, in individuals diagnosed with episodic migraine without aura during the interictal phase, compared with healthy controls.

**Methods:** To that end, 14 migraine patients and 15 healthy controls were recruited (all female), and diffusion-weighted and T1-weighted MRI data were acquired. The structural connectome was estimated for each participant based on two different whole-brain parcellations, including cortical and subcortical regions as well as the cerebellum. The structural connectivity patterns, as well as global and local graph theory metrics, were compared between patients and controls, for each of the two parcellations, using network-based statistics and a generalized linear model (GLM), respectively. We also compared the number of connectome streamlines within specific white matter tracts using a GLM.

**Results:** We found increased structural connectivity in migraine patients relative to healthy controls with a distinct involvement of cerebellar regions, using both parcellations. Specifically, the node degree of the posterior lobe of the cerebellum was greater in patients than in controls and patients presented a higher number of streamlines within the anterior limb of the internal capsule. Moreover, the connectomes of patients exhibited greater global efficiency and shorter characteristic path length, which correlated with the age onset of migraine.

**Conclusions:** A distinctive pattern of heightened structural connectivity and enhanced global efficiency in migraine patients compared to controls was identified, which distinctively involves the cerebellum. These findings provide evidence for increased integration within structural brain networks in migraine and underscore the significance of the cerebellum in migraine pathophysiology.

## 1. Introduction

About 14% of people suffer from migraine, making it one of the most prevalent neurological diseases worldwide [1]. Although its pathophysiology is still not fully understood, multiple studies have found structural [2–11] and functional [2, 10–14] abnormalities throughout extensive brain networks. In particular, the cerebellum has an important role in pain processing, and it has been shown to be altered in numerous neuroimaging studies of migraine [14]. Mehnert et al. [15] reported gray matter volume changes in the crus (I and II), while Qin et al. [16] found microstructural changes in the vermis VI and in the bilateral lobules V and VI of the cerebellum. Evidence also suggests that migraine patients are more prone to ischemic lesions in the posterior lobe of the cerebellum [17]. Importantly, two studies [12, 13] have reported functional connectivity changes involving the crus, as well as the insula and the hippocampus, namely in terms of the network measures of centrality and efficiency. Taken together, these studies highlight the significant role of the cerebellum in the structural and functional brain changes observed in patients with episodic migraine.

Regarding structural connectivity, several studies have reported alterations in migraine based on diffusion Magnetic Resonance Imaging (dMRI) parameters [4, 5, 8–10, 18]. Some studies found increased structural connectivity between cortical regions involved in pain perception and processing [8], and between subcortical regions such as the thalamus and the caudate nucleus [4], suggesting that subcortical networks are strengthened in migraine [4, 5]. Other studies investigated connectome differences from a graph theory perspective using the connectivity between regions as edges and the regions as nodes. Li et. al. [10] found a decreased characteristic path length in the structural connectome of migraine patients, which points to an increased number of short-distance connections and hence an increased integration. Similarly, increased integration was found in subcortical regions such as the putamen, the pallidum, and the thalamus as well as in the parahippocampal gyrus, and the anterior cingulate gyrus [3, 10]. Although the cerebellum has an important role in pain processing, and it has been shown to be altered in numerous neuroimaging studies of migraine [14], to our knowledge no study of structural connectivity using tractography in migraine has included it.

Here, we aim to investigate structural connectivity changes in migraine, considering the whole brain including cortical and subcortical regions as well as the cerebellum. It is important to note that some of the studies mentioned above have heterogeneous cohorts, including male and female patients, with and without aura, and with chronic and episodic migraine [4–6, 15]. This makes it difficult to disentangle the specific connectivity patterns for each subtype of migraine, as it is very likely that different subtypes have different brain functional and connectivity patterns. In our study, we focused solely on low-frequency episodic migraine without aura in female patients, which is by far the most prevalent group of migraine patients [19, 20].

## 2. Methods

### 2.1. Participants

This study analyzed data from the MigN2treat cohort (https://welcome.isr.tecnico.ulisboa.pt/projects/multimodal-neuroimaging-biomarkers-throughout-the-migraine-cycle-towards-neurofeedback-training-for-personalized-anti-migraine-treatment/) cohort which comprised of 14 female patients diagnosed with episodic migraine without aura according to the criteria of the 3rd edition of the International Classification of Headache Disorders (ICHD-III) [21], and 15 healthy controls. Both healthy controls and patients add the following inclusion criteria: a) age between 18 and 55 years; b) had at least 9 years of education; c) had Portuguese as their first language; and the following exclusion criteria: a) diagnosis of a neurological condition (other than migraine for patients); b) diagnosis of any psychiatric disorder (severe anxiety or depressive symptoms were excluded by the use of standardized self-report scales State-Trait Anxiety Inventory and the Zung Self-Rating Depression scale); c) daily use of psychoactive medication including migraine prophylaxis for the patient group; d) being pregnant, breastfeeding, post-menopause, or using of contraception precluding cyclic menses; e) contraindications for performing an MRI (e.g. being claustrophobic); and f) evidence of incident brain lesion or structural abnormalities on the T1-weighted MRI.

Healthy controls were matched for gender, contraceptive use, and menstrual phase at the time of scanning and balanced for age. All participants provided written informed consent and the study was carried out following the Declaration of Helsinki upon approval by the local Ethics Committee.

Imaging data was acquired during the post-ovulation period, corresponding to the interictal phase, as participants in the cohort experienced menstrual-related migraine attacks. Prior to scanning, participants were required to be free from pain for at least 48 hours, with confirmation of the absence of a migraine attack obtained 72 hours post-scan.

The following demographic and clinical parameters were collected: age (years), frequency (migraine attacks per month), age of onset of migraine (years), and disease duration (years).

### 2.2. Data Acquisition

The MRI data was acquired using a 3 Tesla system (Siemens Vida), equipped with a 64-channel receiver head coil from June 2019 to November 2022. Regarding the dMRI data, an echo planar imaging (EPI) sequence was used with the following parameters: Time of Repetition (TR) = 6800 ms, Time to Echo (TE) = 89 ms, 66 axial slices with an in-plane Generalized Autocalibrating Partially Parallel Acquisitions (GRAPPA) factor of 2, Simultaneous Multi-slice (SMS) factor of 3, flip angle = 90°, and 2 mm isotropic voxel size. The sampling scheme consisted of 3 diffusion shells (multi-shell) with b = 400, 1000, 2000 s/mm^2^ along 32, 32, 60 unique gradient directions, respectively, 8 non-diffusion-weighted volumes (b0) and an additional 3 b0 images with opposite phase encoding (Posterior-Anterior). The total imaging time was 15 minutes and 47 seconds. One T1-weighted image was for each subject also acquired with a 3D magnetization-prepared rapid gradient echo (MPRAGE) sequence with the following parameters: 1mm isotropic voxel size, TR = 2300 ms, TE = 2.98 ms, Time of Inversion (TI) = 900 ms, flip angle = 9°, FOV = 256 × 240 × 128 mm^3^ having lasted 5 minutes and 12 seconds. A Fluid-Attenuated Inversion Recovery (FLAIR) image was also acquired and accessed for white matter lesions using the following parameters: 0.9mm isotropic resolution, TR = 5000 ms, TE = 386 ms, TI = 1800 ms, (FOV) = 240 × 240 × 151 mm^3^, GRAPPA = 2, and lasted 5 minutes and 57 seconds, with 0.9mm isotropic resolution. Both anatomical scans were evaluated by an experienced neurologist.

### 2.3. Data Preprocessing

An overview of the image processing and subsequent analyses can be found in Figure 1.

**Figure 1:**
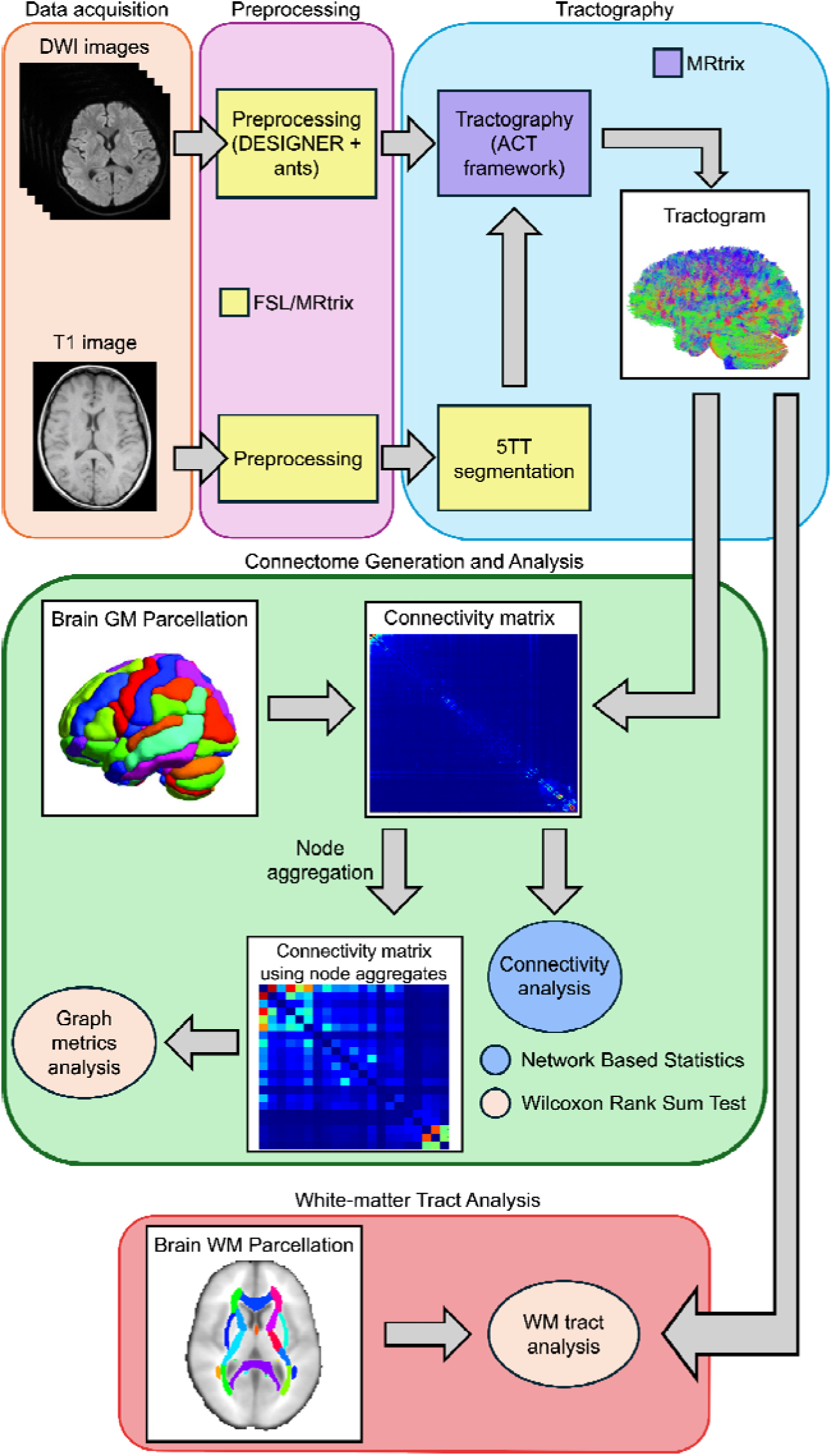
Overview of the data processing pipeline, from the acquisition of the images to the connectome generation, including the different analyses performed.

Image processing was conducted using a combination of FSL [22] (version 6.0.5) and MRtrix [23] (version 3.0.3) tools. The following preprocessing procedures were applied to the dMRI data based on the DESIGNER (Diffusion parameter EStImation with Gibbs and NoisE Removal) pipeline [24]: denoising (*dwidenoise*), Gibbs ringing correction (*mrdegibbs*), and Rician bias correction were performed according to [25], followed by geometric and eddy-current distortions and motion correction with FSL (*eddy* and *topup*). After this, bias field correction was done with MRtrix (*dwibiascorrect* using the -*ants* option).

### 2.4. Tractography

To calculate the streamlines (tracts) of white matter fibers, MRtrix was used. Firstly, the basis functions for each tissue type were estimated from the subject’s dMRI data. Then, the fiber orientation density functions (FODf) were calculated using multi-shell multi-tissue constrained spherical deconvolution [26] followed by their normalization to enable the comparison between subjects. Afterward, Anatomically-Constrained Tractography (ACT) [27] was performed which makes use of prior anatomical knowledge to restrict the generated streamlines by, for instance, preventing streamlines from ending in implausible places such as in the middle of white matter or in the cerebrospinal fluid (CSF). This is done by segmenting the brain into five tissue types (5TT) using the *5ttgen* function of MRtrix and then using the grey and white matter masks to create a seed region along the interface of these two regions from which seed points were randomly generated. From these points, the streamlines were reconstructed thus creating a tractogram. Ten million streamlines were seeded with a maximum length of 25 cm using the iFOD2 tracking algorithm (probabilistic) [28]. Subsequently, spherical deconvolution-informed filtering of tractograms (SIFT2) [29] was performed. Finally, the tractogram was utilized to determine the connectivity between different regions of interest (ROI) and to generate a connectome.

### 2.5. Connectome generation

Two sets of ROIs covering the whole brain were defined using two different parcellations: i) *Schaefer + SC + CB parcellation:* 100 cortical regions from the Schaefer atlas [30] combined with 12 subcortical regions (SC) and 26 cerebellum (CB) regions from the Automated Anatomical Labeling atlas with 116 nodes (AAL116) [31]; and ii) *AAL116 parcellation:* the 116 regions of the AAL116 atlas, which includes cortical regions as well as the same SC and CB regions as in i).

For each of these parcellations, a connectivity matrix was calculated based on the streamlines that connect each pair of ROIs and the volumes of those ROIs, according to:

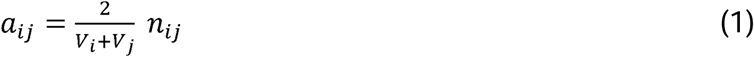

where *a_ij_* is the entry of the connectivity matrix concerning ROI *i* and ROI *j*, *n_ij_* is the number of streamlines between the two ROIs given by the tractogram, and *V_i_* and *V_j_* are the volumes of ROI *i* and ROI j, respectively. Note that, due to the absence of directionality in dMRI, the connectivity matrix is symmetric. Also, self-connections (streamlines that begin and end in the same ROI) were not considered, and as such the connectivity matrix has a zero diagonal.

### 2.6. Connectome Analysis

The analyses present in this section were all performed using MATLAB version: 9.14.0 (R2023a).

#### 2.6.1. Connectivity Analysis

To detect significant connectivity changes between patients and controls, the network-based statistics toolbox (NBS version 1.2) [32] was used, which performs mass-univariate testing at every connection evaluating the null hypothesis at the level of interconnected subnetworks instead of individual connections. It leverages permutation testing to assess the statistical significance across subnetworks of connections. The default value of 3.1 was used as the test statistic’s lower threshold to apply to the network so that only the more significant connections survive. NBS then examines whether these significant connections form larger clusters/subnetworks that are unlikely to have occurred by chance by performing a 5000 permutations (default value) testing. Clusters whose family-wise error rate (FWER) corrected p-value is lower than 0.05 were considered statistically significant. Age was used as a covariate.

For a more straightforward interpretation of the results, the regions (nodes) of each parcellation were grouped into larger regions (node aggregates) according to the larger-scale organization of each atlas: i) *Schaefer + SC + CB parcellation:* the 7 canonical resting-state functional networks identified by Yeo [33] + subcortical regions + crus + posterior lobe of the cerebellum (PLC) + vermis; and ii) *AAL116 parcellation*: 4 lobes (occipital, parietal, frontal and temporal) + subcortical regions + crus + PLC + vermis. For all regions but the vermis, left and right regions were considered separately.

#### 2.6.2. Graph Metrics Analysis

The following graph metrics were calculated on the node aggregates described above using the Brain Connectivity Toolbox (version 2019) [34]: global metrics (characteristic path length, global efficiency, average degree, clustering coefficient, and small-worldness) and a local metric (node degree). For the comparisons between patients and controls, a Generalized Linear Model (GLM) was fitted to the data, including group as the main effect and age as a covariate. In the patients, the correlation between the graph metrics and the clinical parameters was calculated using the Spearman correlation coefficient. For both the group comparisons and the correlations, a p-value lower than 0.05 was considered significant for the global metrics, whilst a p-value lower than 0.05 corrected for the number of nodes using Bonferroni correction was considered significant for the nodal metrics (p<0.0024 and p<0.0033, for the *Schaefer + SC + CB parcellation* and *AAL116 parcellation*, respectively).

#### 2.6.3. White Matter Tract Analysis

Additionally, using the tractogram of each subject, the number of reconstructed streamlines was computed within each ROI of a white matter atlas (ICBM-DTI-81 white-matter labels atlas [35]). A GLM was fitted to the data, which included the group as the main effect and age as a covariate. A p-value lower than 0.001 (corrected for the number of ROIs of the atlas using Bonferroni correction) was considered significant.

## 3. Results

### 3.1. Study population and MRI data

Concerning the clinical and demographic data collected about the subjects, Table 1 presents a statistical analysis performed. No significant changes were found between patients and controls concerning their age (p=0.08).

**Table 1:**
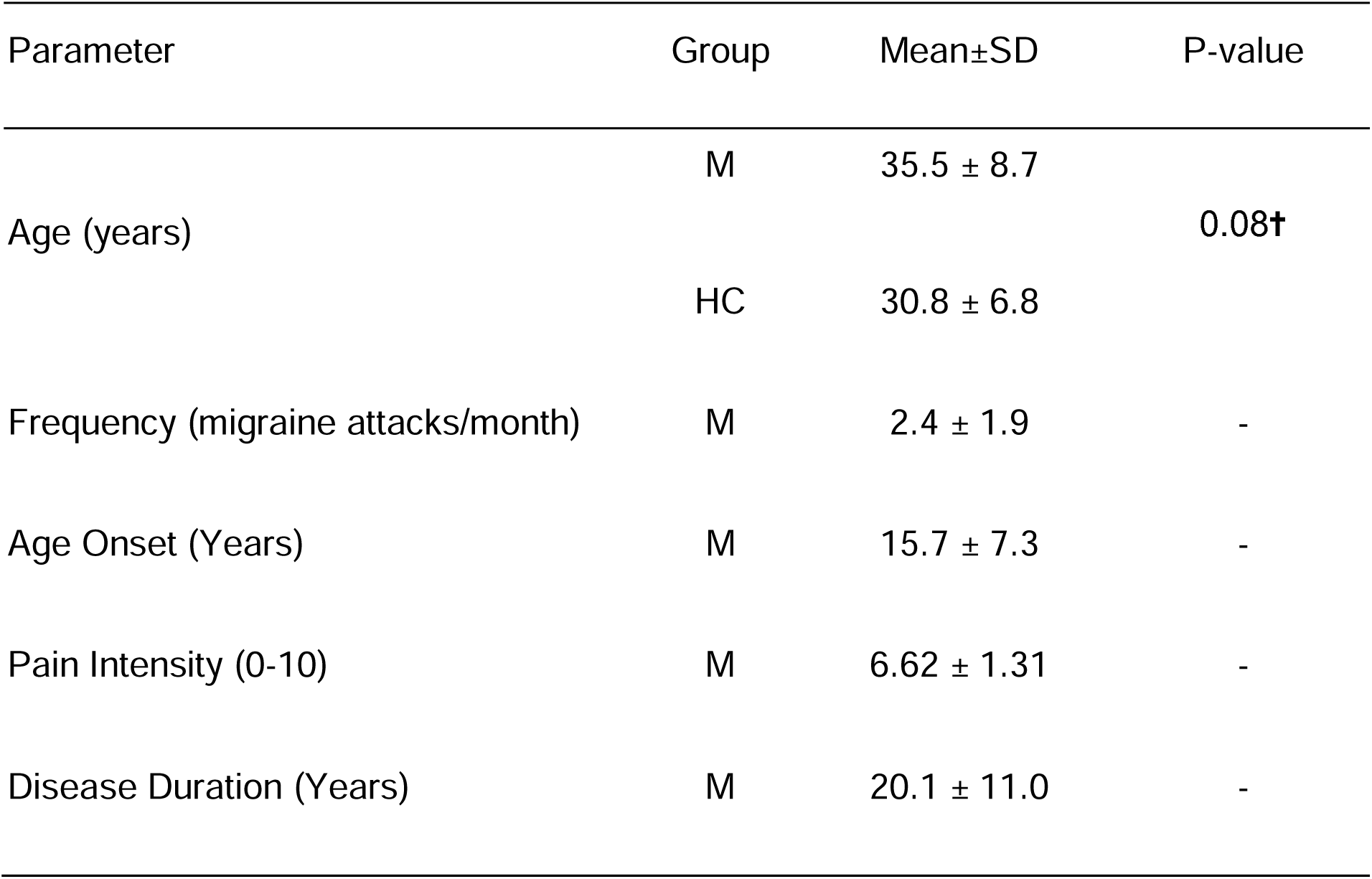
Clinical and demographic characteristics of migraine patients (M) and healthy controls (HC). No significant changes were found between patients and controls concerning their age (p=0.08). †Mann-Whitney U test.

| Parameter | Group | Mean±SD | P-value |
| --- | --- | --- | --- |
| Age (years) | M | 35.5 ± 8.7 | 0.08† |
| | HC | 30.8 ± 6.8 | |
| Frequency (migraine attacks/month) | M | 2.4 ± 1.9 | - |
| Age Onset (Years) | M | 15.7 ± 7.3 | - |
| Pain Intensity (0-10) | M | 6.62 ± 1.31 | - |
| Disease Duration (Years) | M | 20.1 ± 11.0 | - |

Furthermore, no relevant white matter lesions were found in the anatomical scans.

An example of a tractogram and the corresponding connectivity matrices, obtained from a representative patient with each parcellation, are shown in Figure S1 of the Supplementary Material.

### 3.2. Connectivity Analysis

Several connections exhibited significantly increased connectivity in patients relative to controls, for both parcellations (listed in Table S1 and Table S2 in the Supplementary Material). Specifically, a network with increased connectivity in patients was found in both parcellations with p=0.03 (corrected for FWER) for the Schaefer+SC+CB parcellation and p=0.04 (corrected for FWER) for the AAL116 parcellation.

For the visualization of significant connectome differences, we display the total number of connections exhibiting significant differences between patients and controls within each node aggregate in Figure 2, both for the Schaefer + SC + CB parcellation (left) and the AAL116 parcellation (right).

**Figure 2:**
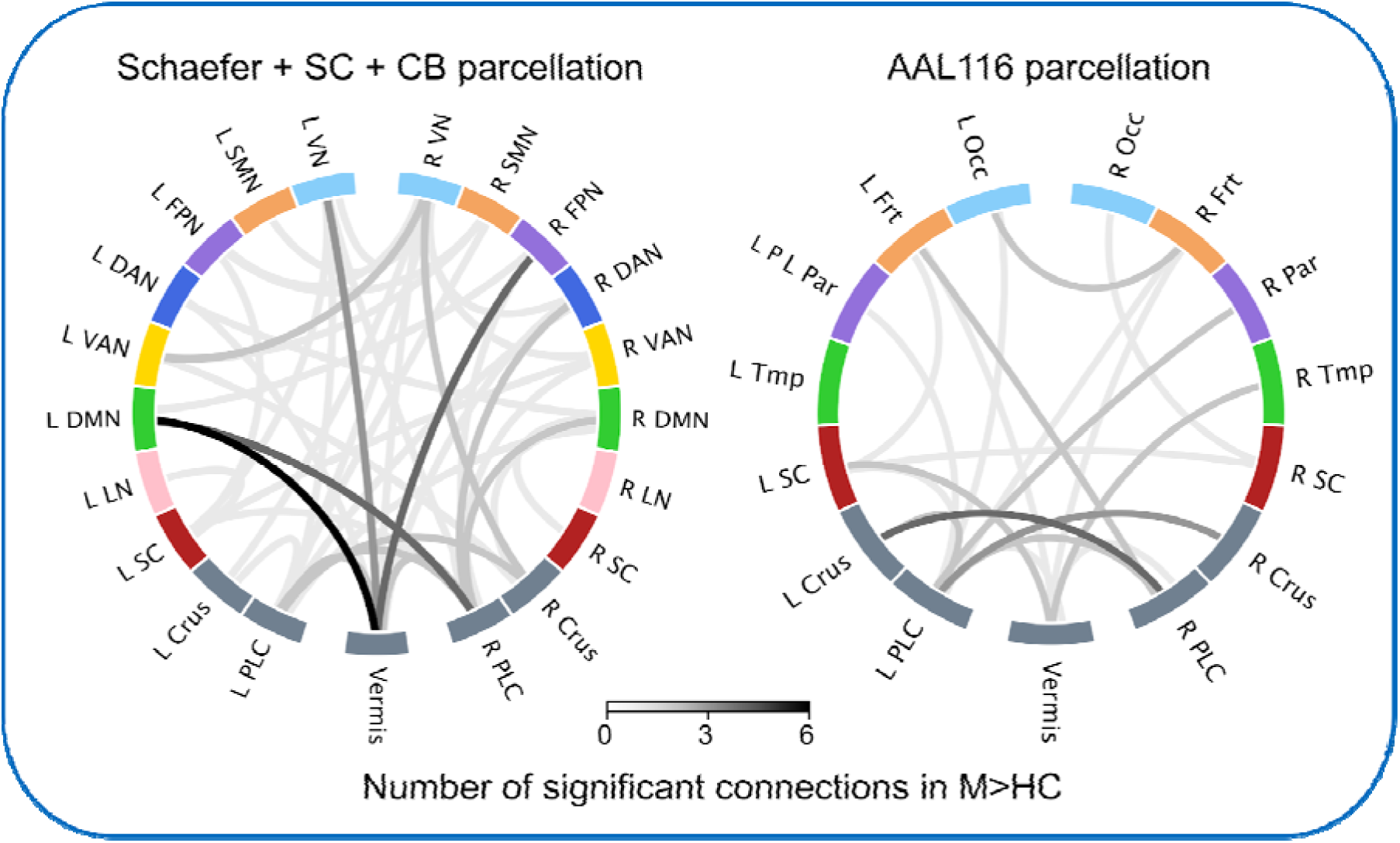
Connectograms of connections exhibiting significantly increased connectivity in patients (M) vs. controls (HC), detected by NBS using both the Schaefer + SC + CB parcellation (M>HC, p=0.03) and the AAL116 parcellation (M>HC, p=0.04). The edges represent the sum of connections with significant differences within each region (node aggregate). L=Left, R=Right, VN=Visual Network, SMN=Somatossensory Network, FPN=Fronto-Parietal Network, DAN=Dorsal Attention Network, VAN=Ventral Attention Network, DMN=Default Mode Network, LN=Limbic Network, Occ=Occipital, Frt=Frontal, Par=Parietal, Tmp=Temporal, SC=Subcortical, PLC=Posterior Lobe of the Cerebellum.

Despite some differences between the results obtained with the two parcellations, common changes in connectivity patterns exist. Specifically, increased connectivity in patients was consistently observed with both parcellations between the right crus and the left posterior lobe of the cerebellum, as well as between the frontal and parietal lobe network and the vermis. Additionally, increased connectivity was also evident between various cerebellar regions and occipital regions (vision network). In summary, a distinctive pattern of increased cerebellar connectivity was consistently observed in patients relative to controls, using both parcellations.

No network was found to have decreased connectivity in migraine patients.

### 3.3. Graph Metrics Analysis

Concerning local graph metrics, the only significant between-groups difference was found in the node degree of the right and left posterior lobe of the cerebellum, which was increased in patients relative to controls using both parcellations, as shown in Figure 3. The results for the global graph metrics are presented in Figure 4. A significant decrease of the characteristic path length and a significant increase of the global efficiency were observed in patients relative to controls. The average degree was significantly higher in patients compared to controls, using both parcellations, while the clustering coefficient and the small-worldness showed no significant differences between groups. The p-values obtained for each metric are presented in Table S3 of the Supplementary Material.

**Figure 3:**
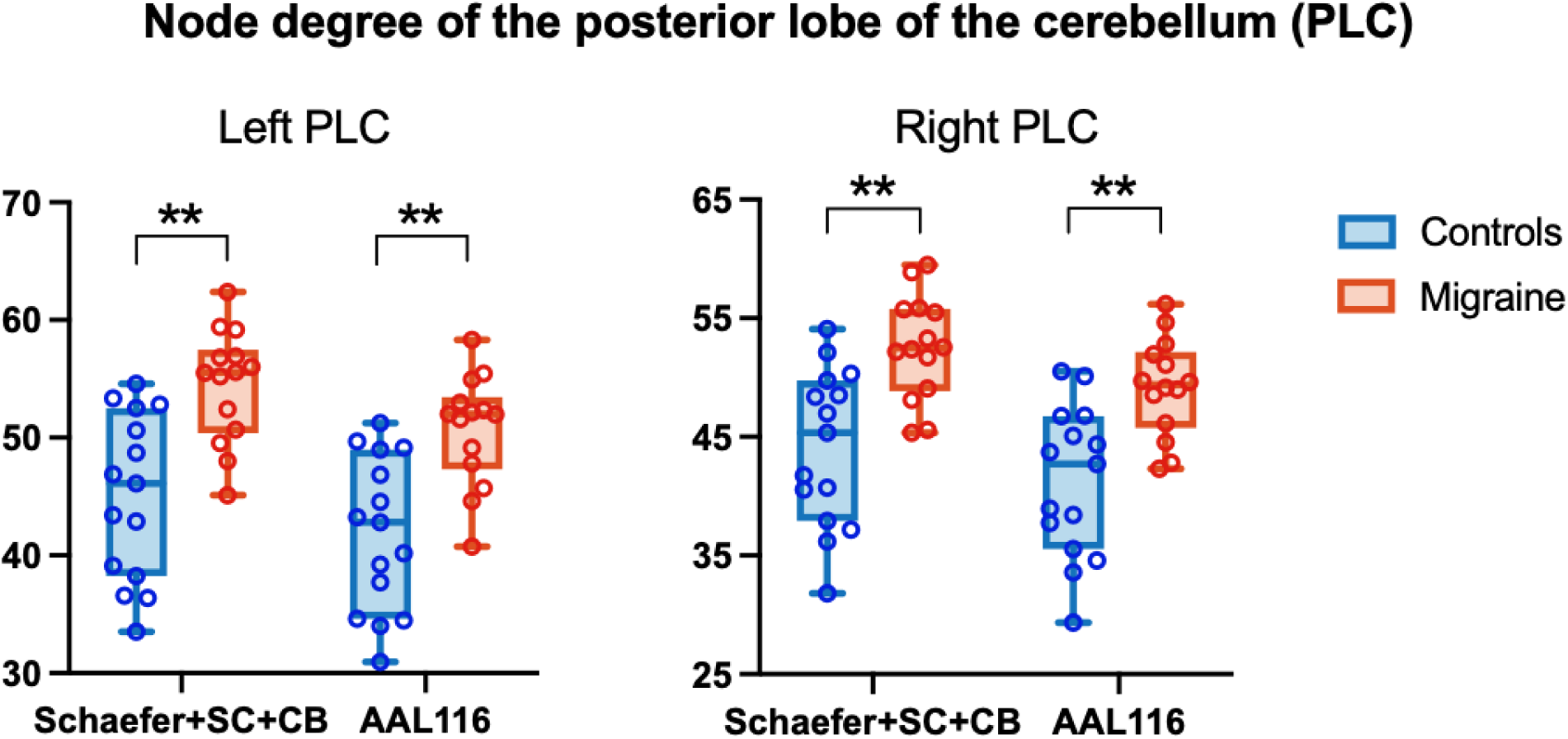
Local graph metrics that show significant differences between patients and controls for both parcellations. The boxplots represent distributions of the metrics across subjects. Significant differences between groups are indicated (*p<0.05, **p<0.01 corrected for multiple comparisons). The node degree of the right and left Posterior Lobes of the Cerebellum (PLC) is increased in migraine patients relative to healthy controls (M>HC).

**Figure 4:**
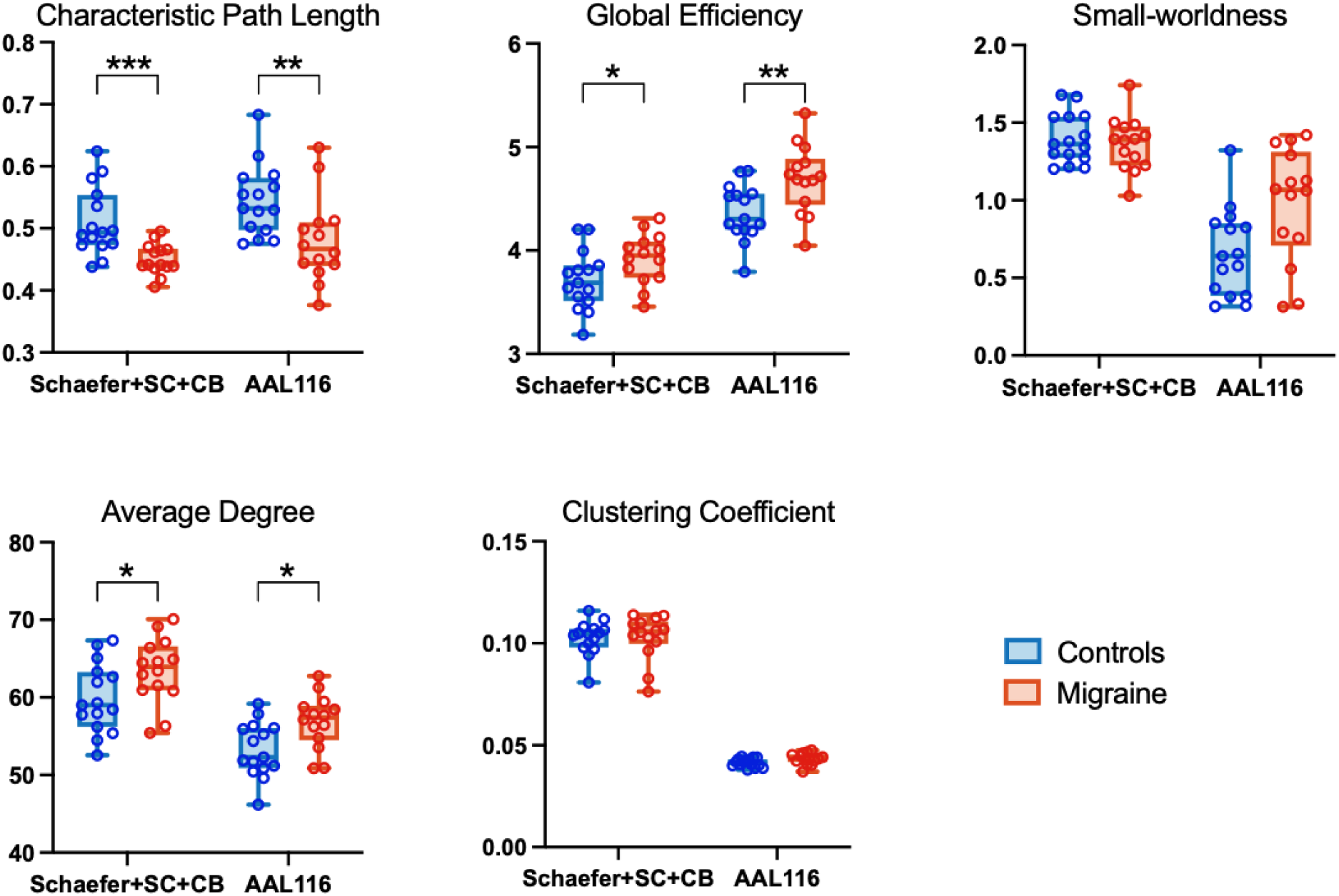
Global graph metrics in patients and controls for both parcellations. The boxplots represent distributions across subjects. Significant differences between groups are indicated (*p<0.05, **p<0.01, ***p<0.001). The characteristic path length was decreased in patients relative to controls in both parcellations, whilst the global efficiency and the average degree were increased. No significant changes were found in the clustering coefficient and the small-worldness.

Moreover, a significant correlation (Spearman rank correlation r^2^=0.26, p=0.038) was found between the characteristic path length obtained using the Schaefer + SC + CB parcellation in patients and their age onset of migraine (Figure 5). No other significant correlations between graph metrics and clinical parameters were found.

**Figure 5:**
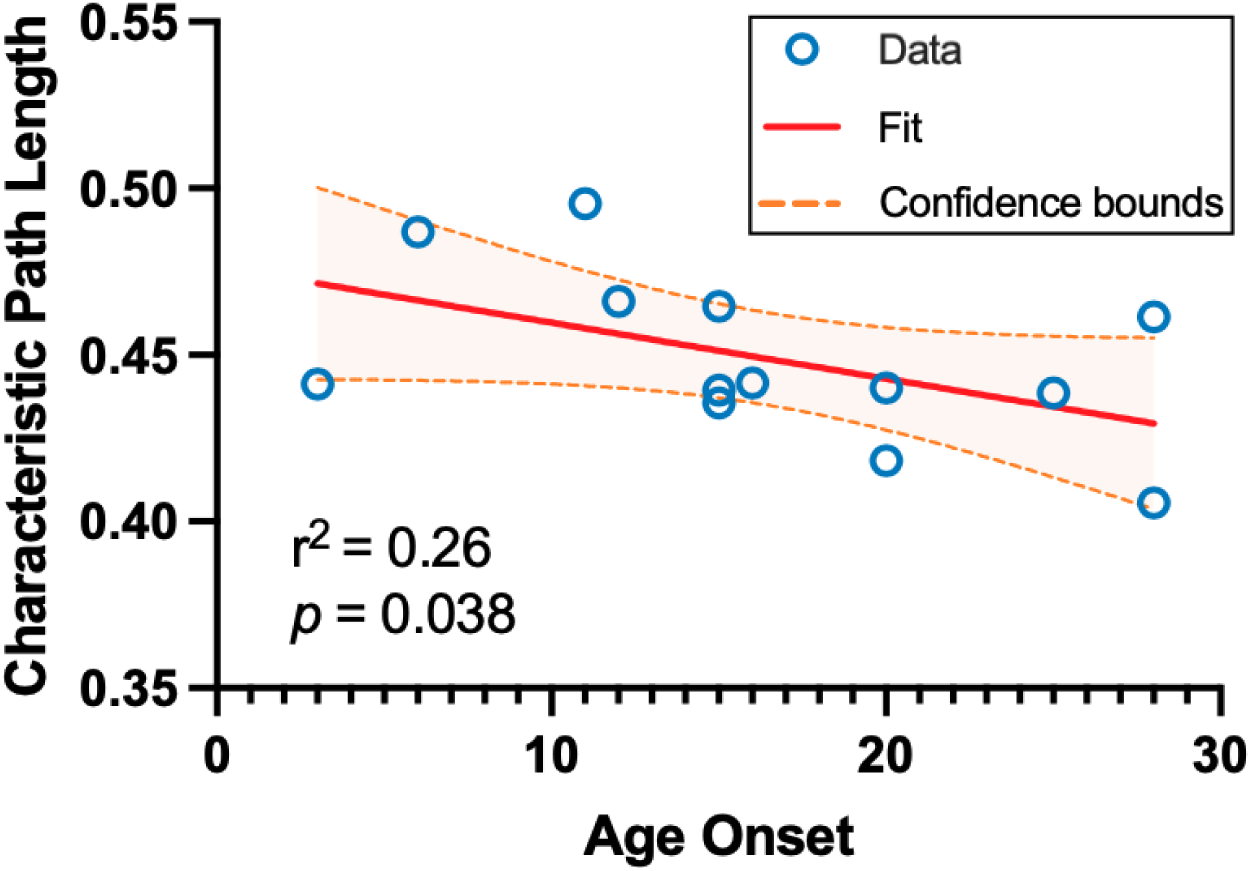
Correlation between the characteristic path length (from the connectome obtained with the Schaefer + SC + CB parcellation) and the age onset of migraine for the patients (Spearman rank correlation r^2^=0.26 with p=0.038).

### 3.4. White matter tract analysis

Regarding the change in the number of reconstructed white matter streamlines, although not statistically significant, there is a slight bilateral increase in the anterior limb of the internal capsule (ALIC) in the patient group relative to controls, as shown in Figure 6.

**Figure 6:**
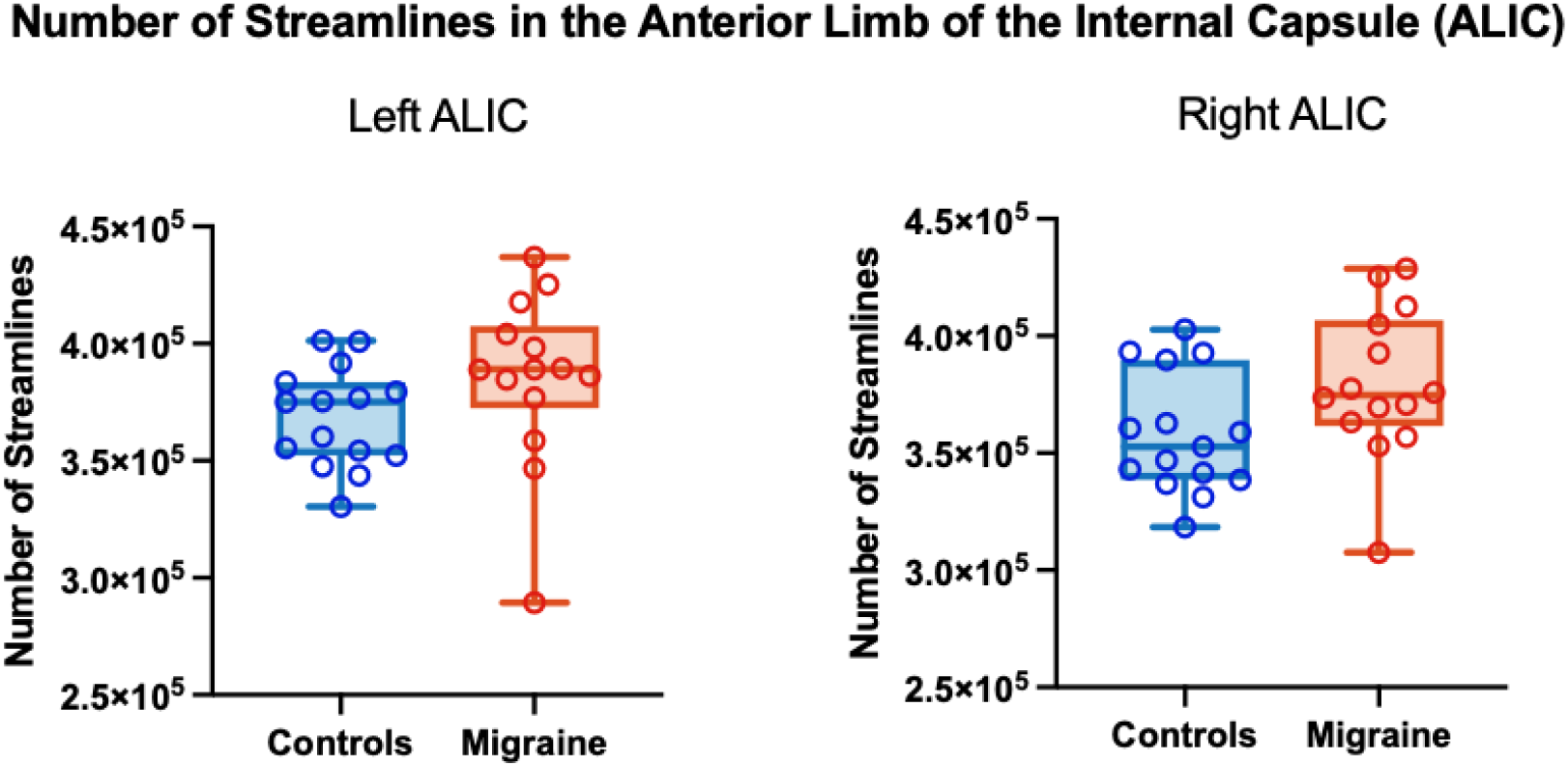
Number of reconstructed streamlines in the left and right anterior limb of the internal capsule (ALIC) (from the ICBM-DTI-81 white-matter labels atlas) for patients (M) and controls (HC). The boxplots represent distributions of the metrics across subjects. No significant differences between groups are indicated. In both ROIs, there was a slight increase in the number of reconstructed streamlines in patients.

## 4. Discussion

In this study, the structural connectome of a group of migraine patients with episodic migraine without aura was studied using dMRI and tractography considering two different whole-brain parcellations. We identified increased connectivity of cerebellar regions in migraine patients, which was reflected in the increased node degree of the posterior lobe of the cerebellum. Consistently with such increased connectivity, we also find an increased number of streamlines in the anterior limb of the internal capsule. From a global connectome perspective, we found increased global efficiency and decreased characteristic path length in patients, which was correlated with the age onset of migraine.

### 4.1. Connectivity analysis

The overall patterns of increased connectivity in patients relative to controls were characterized by a strong involvement of cerebellar regions, namely the crus, the posterior lobe of the cerebellum, and the vermis.

The connectivity analysis showed increased connectivity between the crus and the posterior lobe of the cerebellum. It is believed that the crus exhibits increased activity in response to painful stimuli and has cognitive and emotional representations [36]. Since the crus and the posterior lobe of the cerebellum are involved in cognitive processes [37], alterations in their connectivity may be associated with cognitive impairments reported in migraine patients [38–40]. In addition, stimulation of the posterior lobe of the cerebellum has been shown to modulate the response to noxious visceral stimuli in mice [41], which could be related to migraine pathology.

We also found increased connectivity between the vermis and other regions. The posterior vermis is the anatomical substrate of the limbic cerebellum [37] and thus there can be a disruption in the cerebro-cerebellar-limbic loops of cerebellar input that could be indicative of the emergence of emotional disturbances like mood changes often observed in migraine patients [11, 39].

The cerebellum has an inhibitory role in pain processing [42], having several connections to the prefrontal cortex (via the thalamus) [43]. Therefore, the increased connectivity between the cerebellum and other regions could be indicative of a dysfunctional negative feedback loop in which the thalamus is not sensing the inhibitory signal [42]. These connectivity results could be indicative of the importance of the cerebellum in the modulation of migraine possibly through the modulation of calcitonin gene-related peptide (CGRP) [14, 15]. CGRP modulates nociception and assists in the onset of migraine, and thus it might play a role in reshaping connectivity and stabilizing synapses in the cerebellar circuitry as shown in early preclinical studies [44, 45]. In fact, many migraine-specific high-efficacy treatments target CGRP [46, 47], hence changes in the connectivity pattern in the cerebellum could be potentially explored both as potential biomarkers for treatment response or as a predictor of treatment efficacy.

Our results not only contribute further evidence that the cerebellum plays a central role in migraine pathophysiology but also emphasize the importance of including it as a region of interest in connectome studies of migraine, with the robustness of our analysis being supported by the observation of similar patterns across two different brain parcellations.

### 4.2. Graph metrics analysis

At the global connectome level, decreased characteristic path length and increased global efficiency were found in patients compared to controls, which can be associated with a dysfunctional modulation of pain networks in migraine and a faster dissemination of pain-related information [8–10]. Moreover, the negative correlation found between the characteristic path length of patients and their age onset could potentially imply a plastic adaptation to migraine over time (since the patient developed this disorder), which has been reported by several diffusion tensor imaging studies [4, 6, 48, 49] in migraine patients. These changes could also contribute to an increased pain perception as has been reported in functional studies [50, 51]. Additionally, some studies have also reported correlations between graph theory metrics or diffusion metrics and patients’ clinical data. For instance, Dai et al. [9] reported a correlation between local efficiency and the visual analog scale in episodic migraine without aura, while Planchuelo-Gómez et al. [4] and Chong et al. [6] reported a relationship between diffusion metrics such as fractional anisotropy (FA) and mean diffusivity and the years lived with migraine (disease duration), using less homogeneous cohorts than ours, suggesting that chronicity might be exacerbating neural abnormalities.

In terms of local graph metrics, we found a bilateral increase in the node degree of the posterior lobe of the cerebellum, which is consistent with the increased connectivity of this region with the rest of the brain. As the posterior lobe of the cerebellum has both cognitive and limbic functions [52], the increased node degree of this region might be a result of a compensatory mechanism leading to effects on cognitive processing and emotional state.

### 4.3. White matter tract analysis

Complementarily to the analysis of the structural connections between gray matter regions, the corresponding white matter tracts were also evaluated considering the number of streamlines that passed through each tract and found a slight increase in the number of streamlines bilaterally in the anterior limb of the internal capsule of patients. The internal capsule contains ascending and descending axons including fibers to and from the thalamus, and to and from the cerebellum [53]. This increase is therefore consistent with the increased connectivity of the cerebellum, and could also be indicative of the dysfunctional negative feedback loop mentioned before. On the other hand, the increased number of reconstructed fibers could also be a consequence of an increased FA in the cerebellum of patients, which other researchers [54] have found. Future studies should clarify these different possible reasons.

### 4.4. Limitations and innovative aspects

This study has a few limitations, the main one being the small sample size. Despite that, we focused on a homogeneous patient group with controlled conditions and selected a well-matched control group in terms of gender, and menstrual cycle phase, minimizing individual variability.

Furthermore, our connectivity metrics are based on the number of reconstructed streamlines obtained by using a specific tractography algorithm. Although this methodology is well-established and commonly used in the literature, it comes with several challenges in terms of quantification and interpretation. However, throughout the tractography pipeline, decisions were made to tackle these challenges. For instance, two of the main biases of this type of analysis are streamline termination biases, where the reconstructed streamlines end in unreasonable voxels (such as in the middle of CSF), and streamline quantification biases where the number of reconstructed fibers does not represent the actual fiber count for a given voxel [55, 56]. In this work, we employed the ACT framework that tackles the first bias by seeding the streamlines from the interface between grey and white matter, and we used a streamline filtering technique (SIFT2) to deal with the quantification of fiber density in each voxel. Currently, using the number of streamlines as a proxy for fiber count allied to these advanced post-processing techniques is the state-of-the-art for structural connectivity analysis generation [55]. Finally calculating local and global graph theory metrics provides a complete assessment of structural connectome. By using weighted connectivity matrices to represent the structural connectome instead of binary matrices, there is a better representation of biological properties [57].

This is the first study that uses tractography based on multi-shell data to estimate the structural connectome in migraine. Moreover, the inclusion of the cerebellum in two different brain parcellations is also novel. Finally, our study is unique by focusing on a homogeneous cohort of female-only patients with menstrual and menstrual-related low-frequency episodic migraine, and including healthy controls matched for the menstrual phase.

## 5. Conclusion

In conclusion, our findings reveal structural connectome alterations in a cohort of patients diagnosed with menstruation-related low-frequency episodic migraine without aura, prominently implicating the cerebellum. These results substantiate the notion of heightened integration across whole brain networks underscoring the previously acknowledged yet often overlooked role of the cerebellum in migraine pathophysiology.

## Acknowledgments

We acknowledge the Portuguese Science Foundation and LARSyS for supporting our study. We also express our gratitude to this study’s participants and the collaborators from Hospital da Luz Lisboa.

