## Supplementary Material for "Involvement of the cerebellum in structural connectivity enhancement in episodic migraine"

### Figure S1

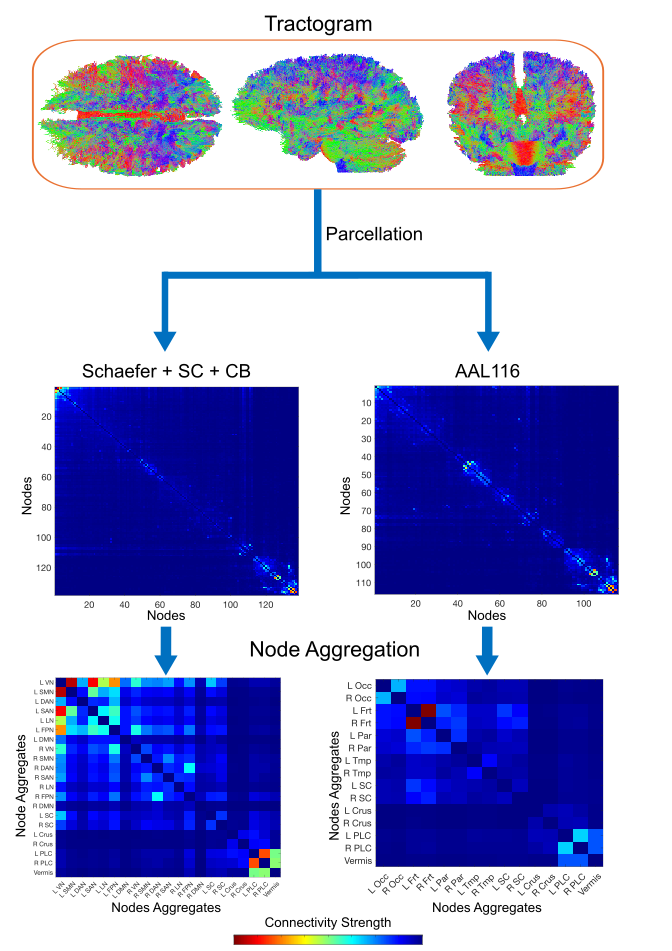

Figure S1: Example of a tractogram and the connectivity matrices obtained with the two parcellations (Schaefer + SC + CB and AAL116) before (top) and after (bottom) node aggregation.

Table S1: Pairs of connectome nodes whose connection NBS detects as being significantly increased in patients when using the Schaefer + SC + CB parcellation (p=0.03).

| Node 1 | Node 2 |
| --- | --- |
| Vis_1-L | Caudate-L |
| Vis_1-L | Cerebelum-9-L |
| Vis_1-L | Vermis-6 |
| Vis_2-L | Vis_8-R |
| Vis_4-L | Vis_8-L |
| Vis_4-L | Cerebelum-9-R |
| Vis_6-L | Vermis-7 |
| Vis_9-L | Vermis-7 |
| SomMot_3-L | Vis_8-R |
| DorsAttn_Post_2-L | Default_Temp_1-R |
| DorsAttn_FEF_1-L | Vermis-7 |
| SalVentAttn_ParOper_1-L | Cerebelum-Crus2-R |
| SalVentAttn_FrOperIns_1-L | Vis_8-R |
| SalVentAttn_FrOperIns_2-L | SomMot_1-R |
| SalVentAttn_Med_3-L | Vis_8-R |
| Limbic_TempPole_1-L | Putamen-L |
| Cont_Cing_1-L | Vis_8-R |
| Cont_Cing_1-L | SomMot_1-R |
| Cont_Cing_1-L | Putamen-L |
| Default_Par_2-L | Cerebelum-3-R |
| Default_PFC_1-L | Vermis-6 |
| Default_PFC_6-L | Cerebelum-3-R |
| Default_PFC_6-L | Vermis-7 |
| Default_PFC_7-L | Cerebelum-8-R |
| Default_PFC_7-L | Cerebelum-9-R |
| Default_pCunPCC_1-L | Vermis-3 |
| Default_pCunPCC_1-L | Vermis-4-5 |
| Default_pCunPCC_1-L | Vermis-6 |
| Default_pCunPCC_1-L | Vermis-7 |
| Default_pCunPCC_2-L | Cont_PFCl_1-R |
| Vis_2-R | Cerebelum-Crus1-R |
| Vis_2-R | Cerebelum-Crus2-L |
| Vis_2-R | Vermis-6 |
| Vis_3-R | DorsAttn_PrCv_1-R |
| Vis_5-R | SalVentAttn_Med_2-R |
| Vis_5-R | Cerebelum-Crus1-R |
| Vis_6-R | Cerebelum-7b-L |
| SomMot_5-R | Cerebelum-9-L |
| DorsAttn_Post_1-R | Cerebelum-3-R |
| DorsAttn_PrCv_1-R | Cerebelum-3-L |
| DorsAttn_PrCv_1-R | Cerebelum-9-R |
| DorsAttn_FEF_1-R | Vermis-3 |
| SalVentAttn_FrOperIns_1-R | Cerebelum-9-R |
| SalVentAttn_Med_2-R | Putamen-L |
| Cont_PFCl_1-R | Cerebelum-3-R |
| Cont_pCun_1-R | Vermis-1-2 |
| Cont_pCun_1-R | Vermis-6 |
| Cont_pCun_1-R | Vermis-7 |
| Cont_pCun_1-R | Vermis-8 |
| Default_Par_1-R | Cerebelum-7b-R |
| Default_Temp_1-R | Pallidum-R |
| Default_Temp_1-R | Cerebelum-Crus2-R |
| Default_PFCdPFCm_1-R | Cerebelum-3-L |
| Default_PFCdPFCm_2-R | Cerebelum-8-R |
| Caudate-L | Cerebelum-7b-R |
| Cerebelum-Crus1-R | Cerebelum-7b-R |
| Cerebelum-Crus2-L | Cerebelum-6-L |
| Cerebelum-Crus2-R | Cerebelum-6-L |
| Cerebelum-Crus2-R | Cerebelum-7b-L |
| Cerebelum-3-L | Cerebelum-4-5-R |
| Cerebelum-3-L | Vermis-3 |
| Cerebelum-4-5-R | Cerebelum-8-R |
| Cerebelum-4-5-R | Cerebelum-9-L |
| Cerebelum-4-5-R | Vermis-3 |
| Cerebelum-6-R | Cerebelum-7b-R |
| Cerebelum-6-R | Vermis-1-2 |
| Cerebelum-9-L | Vermis-6 |

Table S2: Pairs of connectome nodes whose connection NBS detects as being significantly increased in patients when using the AAL116 parcellation (p=0.04).

| Node 1 | Node 2 |
| --- | --- |
| Precentral-L | Cerebelum-9-R |
| Precentral-R | Lingual-L |
| Precentral-R | Vermis-7 |
| Frontal-Sup-L | Pallidum-L |
| Frontal-Sup-L | Cerebelum-9-R |
| Frontal-Sup-Medial-L | Vermis-7 |
| Insula-R | Lingual-L |
| Insula-R | Cerebelum-9-L |
| ParaHippocampal-R | Vermis-4-5 |
| ParaHippocampal-R | Vermis-6 |
| Cuneus-R | Putamen-R |
| Occipital-Inf-L | Cerebelum-9-L |
| Parietal-Sup-R | Cerebelum-9-L |
| SupraMarginal-L | Cerebelum-9-L |
| SupraMarginal-R | Cerebelum-9-L |
| Caudate-L | Cerebelum-9-L |
| Caudate-L | Vermis-6 |
| Putamen-R | Pallidum-L |
| Pallidum-L | Vermis-3 |
| Cerebelum-Crus1-L | Cerebelum-9-R |
| Cerebelum-Crus1-R | Cerebelum-8-L |
| Cerebelum-Crus1-R | Cerebelum-9-L |
| Cerebelum-Crus2-L | Cerebelum-3-L |
| Cerebelum-Crus2-L | Cerebelum-3-R |
| Cerebelum-Crus2-L | Cerebelum-4-5-R |
| Cerebelum-Crus2-L | Cerebelum-6-L |
| Cerebelum-Crus2-L | Cerebelum-9-R |
| Cerebelum-Crus2-R | Cerebelum-9-L |
| Cerebelum-3-L | Vermis-3 |
| Cerebelum-4-5-R | Vermis-3 |
| Cerebelum-8-L | Cerebelum-9-L |
| Cerebelum-8-R | Cerebelum-9-L |
| Cerebelum-9-L | Cerebelum-9-R |

Table S3: Significance of the differences between groups for each graph metric. In bold there are the statistically significant differences. L=Left, R=Right, VN=Visual Network, SMN=Somatosensory Network, FPN=Fronto-Parietal Network, DAN=Dorsal Attention Network, VAN=Ventral Attention Network, DMN=Default Mode Network, LN=Limbic Network, Occ=Occipital, Frt=Frontal, Par=Parietal, Tmp=Temporal, SC=Subcortical PLC=Posterior Lobe of the Cerebellum.

| Parcellation | | Schaefer+SC+CB | | AAL116 | |
| --- | --- | --- | --- | --- | --- |
| Local Metric | Degree | Region | p-value | Region | p-value |
|  |  | L VN | 0.5338 | L Occ | 0.5487 |
|  |  | L SMN | 0.5222 | R Occ | 0.6501 |
|  |  | L DAN | 0.4464 | L Frt | 0.1171 |
|  |  | L VAN | 0.5817 | R Frt | 0.1165 |
|  |  | L LN | 0.2295 | L Par | 0.4812 |
|  |  | L FPN | 0.2720 | R Par | 0.7028 |
|  |  | L DMN | 0.2106 | L Temp | 0.0832 |
|  |  | R VN | 0.1065 | R Temp | 0.1212 |
|  |  | R SMN | 0.4478 | L SC | 0.1244 |
|  |  | R DAN | 0.2664 | R SC | 0.0652 |
|  |  | R VAN | 0.4882 | L Crus | 0.0026 |
|  |  | R LN | 0.0995 | R Crus | 0.0058 |
|  |  | R FPN | 0.1088 | L PLC | **0.0002** |
|  |  | R DMN | 0.0601 | R PLC | **0.0002** |
|  |  | L SC | 0.1002 | Vermis | 0.0264 |
|  |  | R SC | 0.1000 |  | |
|  |  | L Crus | 0.0033 |  |  |
|  |  | R Crus | 0.0079 |  |  |
|  |  | L PLC | **0.0001** |  |  |
|  |  | R PLC | **0.0002** |  |  |
|  |  | Vermis | 0.0204 |  |  |
| Global Metrics | Characteristic Path Length | - | **0.0077** | - | **0.0008** |
|  | Global Efficiency | - | **0.0078** | - | **0.0427** |
|  | Clustering Coefficient | - | 0.1146 | - | 0.7409 |
|  | Average Degree | - | **0.0117** | - | **0.0466** |
|  | Small-worldness | - | 0.1501 | - | 0.0977 |
